# Trained-feature specific offline learning in an orientation detection task

**DOI:** 10.1101/429761

**Authors:** Masako Tamaki, Zhiyan Wang, Takeo Watanabe, Yuka Sasaki

**Affiliations:** Department of Cognitive, Linguistic, and Psychological Sciences, Brown University, Box 1821, 190 Thayer Street, Providence, RI, 02912, USA.

## Abstract

It has been suggested that sleep provides additional enhancement of visual perceptual learning (VPL) acquired before sleep, termed offline performance gains. A majority of the studies that found offline performance gains of VPL used discrimination tasks including the texture discrimination task (TDT). This makes it questionable whether offline performance gains on VPL are generalized to other visual tasks. The present study examined whether a Gabor orientation detection task, which is a standard task in VPL, shows offline performance gains. In Experiment 1, we investigated whether sleep leads to offline performance gains on the task. Subjects were trained with the Gabor orientation detection task, and re-tested it after a 12-hr interval that included either nightly sleep or only wakefulness. We found that performance on the task improved to a significantly greater degree after the interval that included sleep and wakefulness than the interval including wakefulness alone. In addition, offline performance gains were specific to the trained orientation. In Experiment 2, we tested whether offline performance gains occur by a nap. Also, we tested whether spontaneous sigma activity in early visual areas during non-rapid eye movement (NREM) sleep, previously implicated in offline performance gains of TDT, was associated with offline performance gains of the task. A different group of subjects had a nap with polysomnography. The subjects were trained with the task before the nap and re-tested after the nap. The performance of the task improved significantly after the nap only on the trained orientation. Sigma activity in the trained region of early visual areas during NREM sleep was significantly larger than in the untrained region, in correlation with offline performance gains. These aspects were also found with VPL of TDT. The results of the present study demonstrate that offline performance gains are not specific to a discrimination task such as TDT, and can be generalized to other forms of VPL tasks, along with trained-feature specificity. Moreover, the present results also suggest that sigma activity in the trained region of early visual areas plays an important role in offline performance gains of VPL of detection as well as discrimination tasks.

## Introduction

After the initial acquisition of a skill, a learning state goes through an offline process, through which further improvement on performance is achieved without actual training (Karni et al., 1998; Walker, 2005), termed offline performance gains. It has been suggested that sleep plays an essential role in offline performance gains in various types of learning (Gais et al., 2000; Maquet et al., 2000; Stickgold et al., 2000a; Stickgold et al., 2000b; Laureys et al., 2001; Gais et al., 2002; Walker et al., 2002; Mednick et al., 2003; Huber et al., 2004; Stickgold, 2005; Walker, 2005; Walker and Stickgold, 2005; Walker et al., 2005; Matarazzo et al., 2008; Tamaki et al., 2008; Yotsumoto et al., 2009b; Diekelmann and Born, 2010; Born and Wilhelm, 2012; Mascetti et al., 2013; Rasch and Born, 2013; Tamaki et al., 2013; Bang et al., 2014; McDevitt et al., 2014; Tononi and Cirelli, 2014). One type of learning for which sleep is beneficial is visual perceptual learning (VPL) (Karni et al., 1994; Gais et al., 2000; Stickgold et al., 2000a; Mednick et al., 2003; Censor et al., 2006; Yotsumoto et al., 2009b; Bang et al., 2014). VPL refers to a long-lasting performance improvement on a visual task after visual experience (Sasaki et al., 2010; Lu et al., 2011; Sagi, 2011), and has been proposed to primarily involve reorganization in early visual areas (Schoups et al., 2001; Schwartz et al., 2002; Shibata et al., 2017) (but see other studies (Law and Gold, 2008; Xiao et al., 2008)). After sleep, either night sleep or a daytime nap, the performance on VPL is significantly enhanced compared to that before sleep. In addition, the amount of improvement after sleep surpasses that after the same amount of time, which does not include sleep (Karni et al., 1994; Gais et al., 2000; Stickgold et al., 2000a; Mednick et al., 2003; Censor et al., 2006; Yotsumoto et al., 2009b; Bang et al., 2014).

However, it remains to be elucidated whether offline performance gains are generalized to VPL tasks. The majority of studies which found that sleep plays a role in the performance gain in VPL used the texture discrimination task (or ‘TDT’), which is a standard task in VPL (Karni et al., 1994; Gais et al., 2000; Stickgold et al., 2000a; Mednick et al., 2003; Censor et al., 2006; Yotsumoto et al., 2009b; Bang et al., 2014). Curiously, other studies that found offline performance gains of sleep in VPL also used a discrimination task, including coarse orientation discrimination (Matarazzo et al., 2008; Mascetti et al., 2013) and motion direction discrimination tasks (McDevitt et al., 2014). Importantly, it has been suggested that the underlying mechanisms between a detection task and a discrimination task are significantly different (Regan and Beverley, 1985; Jazayeri and Movshon, 2006; Bridwell et al., 2013). This raises the question as to whether sleep is effective on VPL in general, including a detection task, or whether the effect of sleep on VPL is specific to discrimination tasks.

In addition, it is controversial which spontaneous brain activity during sleep is involved in offline performance gains of VPL. Sleep spindles (13–16 Hz) are characteristic spontaneous brain activity during non-rapid eye movement (NREM) sleep typically used for sleep stage scoring and appear around the central brain regions. Sleep spindles have been shown to be involved in motor memory in human (Manoach and Stickgold, 2009; Manoach et al., 2010; Tamaki et al., 2013; Manoach et al., 2016; Laventure et al., 2018) and have been suggested to enhance plasticity (Sejnowski and Destexhe, 2000; Rosanova and Ulrich, 2005). Sigma activity (13–16 Hz) is a spontaneous oscillatory activity during sleep whose frequency corresponds to sleep spindles. Since VPL is assumed to involve early visual areas, instead of typical sleep spindles appearing around the central brain regions, previously we measured the strength of regional sigma activity from early visual areas during sleep and found the correlation with the degree of offline performance gains of TDT (Bang et al., 2014). In the current study, we tested whether regional sigma activity in early visual areas is also involved in offline performance gains of the Gabor orientation detection task.

First, the results show that the performance on the Gabor orientation detection task improved significantly after the night of sleep without any additional training. No such offline performance gain was found after the same amount of interval that included only wakefulness. Second, we found that offline performance gains occur after a daytime nap as well. Moreover, sigma activity was significantly larger in the trained region than in the untrained region in early visual areas. The performance improvement over the nap significantly correlated with the power of sigma activity in the trained region of early visual areas. The present results suggest that offline performance gains occur not only a discrimination task but also in a detection task, and that sigma activity in the trained region of early visual areas during sleep may play a common role in offline performance gains of VPL.

## Material and Methods

### Participants

We conducted a careful screening process for eligibility for participation, since various factors are known to influence visual sensitivity and sleep structures. All the subjects had no prior experience in VPL tasks, as experiences in VPL tasks may cause a long-term visual sensitivity change (Karni and Sagi, 1991; Schwartz et al., 2002; Seitz et al., 2005; Yotsumoto et al., 2009a; Sasaki et al., 2010; Lu et al., 2011; Sagi, 2011). People who play action video games frequently were excluded because extensive video game playing affects visual and attention processing (Green and Bavelier, 2003; Li et al., 2009; Berard et al., 2015). In addition, subjects were required to have a regular sleep schedule and anyone who had a physical or psychiatric disease, was currently under medication, or was suspected to have a sleep disorder was excluded, as these factors are known to impact sleep structures (Horikawa et al., 2013; Tamaki et al., 2014; 2016). None had medical conditions, including sleep disorders according to the self-reports. All subjects gave written informed consent for their participation in experiments. This study was approved at the institutional review board at Brown University.

A total of seventeen young healthy subjects with normal or corrected-to-normal vision participated in the study. A total of eight subjects participated in Experiment 1. They were randomly assigned to the sleep group (four subjects; 3 females, 20.0 ± 0.48 years old, mean ± SEM) or to the wake group (four subjects; 3 females, 23.0 ± 1.22 years old, mean ± SEM). A total of nine subjects (4 females, 24.3 ± 0.55 years old, mean ± SEM) participated in Experiment 2. In Experiment 2, which was a within-subject design, all subjects had a nap.

### Experimental design for Experiment 1

The subjects in the sleep group arrived at the experimental room at 9 pm. We explained how to perform the task (the introductory session, see below). Then the subjects performed a pre-training test session of the orientation detection task (see ***Orientation detection task*** below) to measure their initial performance level before training (**Fig. 1A, Sleep group**). After the pre-training test session, which took approximately 5 min, the subjects underwent intensive training on the task for approximately 45 min. After the training session, a post-training test session (approximately 5 min) was conducted to measure the performance change by training. These 3 sessions (pre-training test, training, post-training test) lasted about one hour in total to complete. There were approximately 2-min breaks between the pre-training test and the onset of training sessions, as well as between the training and the onset of post-training test sessions. After the completion of the post-training test session, the subjects slept at their home. In the next morning (9am), the subjects performed a post-interval test session. The post-interval test session lasted approximately 5 min.

**Fig. 1.**
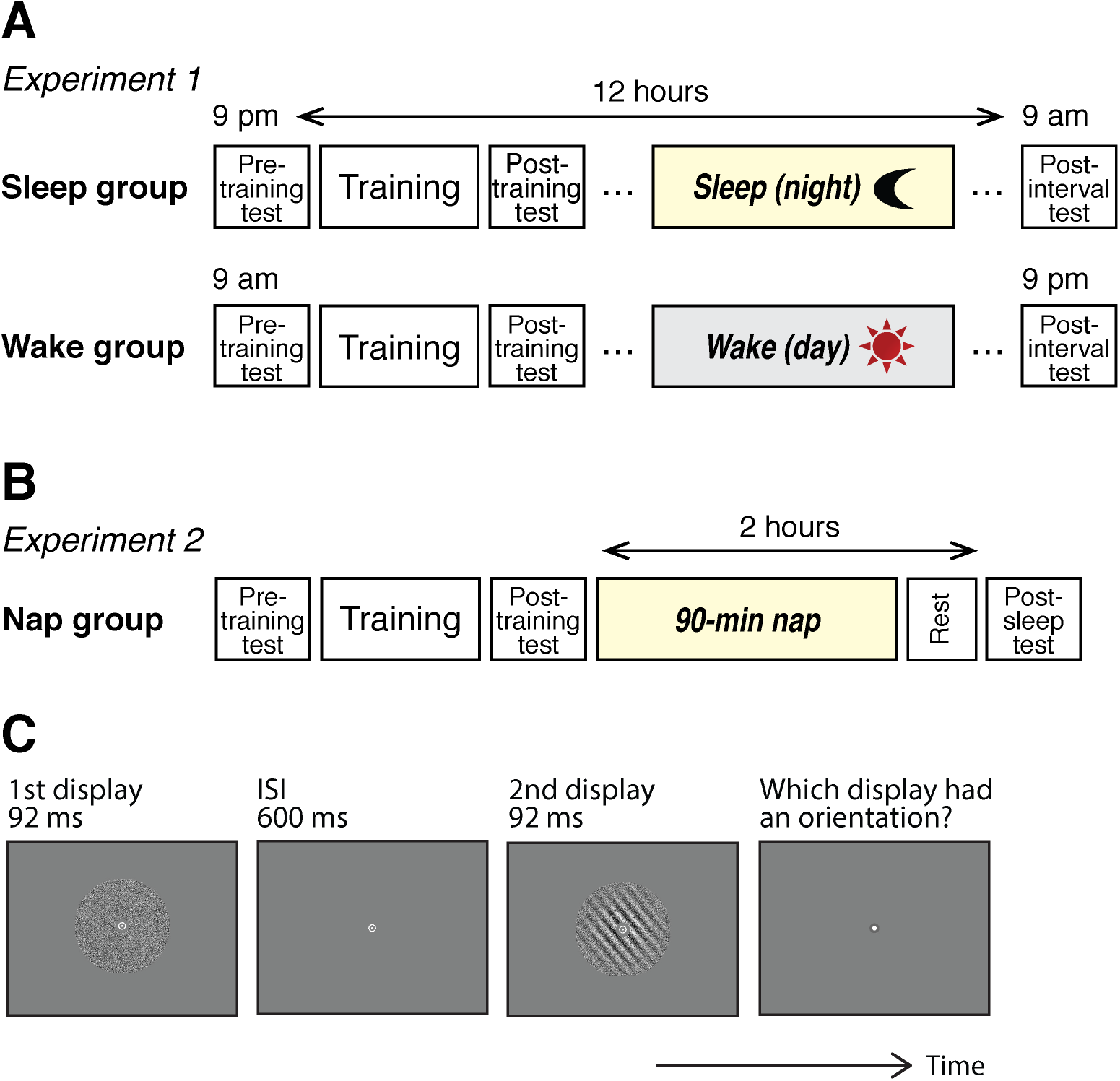
Experimental Designs and the Gabor orientation task. (**A**) Experiment 1. (**B**) Experiment 2. (**C**) Schematic illustration of one trial of the Gabor orientation task.

The subjects in the control wake group arrived at the experimental room at 9 am. After the introduction session, they performed a pre-training test session, training session and a post-training test session, with 2-min break in-between. These sessions took place in the morning between 9–10am (**Fig. 1A, Wake group**). At 9 pm on the same day, they performed a post-interval test session. No sleep was allowed during the day for the Wake group.

The degree of subjective sleepiness was measured by the Stanford Subjective Sleepiness (SSS) scale (Hoddes et al., 1973) before each of the test sessions for both groups.

### Experimental design for Experiment 2

Subjects in Experiment 2 underwent two sleep sessions at our sleep laboratory, adaptation and main experimental sleep sessions. The adaptation session was conducted prior to the main experimental sleep session for the following reason. It has been shown that when subjects sleep in a sleep laboratory for the first time, the sleep quality is degraded due to the new environment, termed the first-night effect (FNE) (Agnew et al., 1966; Carskadon and Dement, 1981; Tamaki et al., 2005; Tamaki et al., 2016). To mitigate the FNE, an adaptation sleep session was necessary prior to the main experimental sleep session. During the adaptation sleep session, all electrodes were attached to the subjects for polysomnography (PSG) measurement. The subjects slept in the same fashion as in the experimental sleep session. The adaptation session was conducted approximately a week before the main experimental sleep session so that any effects due to sleeping during the adaptation sleep session would not carry over to the experimental sleep session.

On the day of the main experimental sleep session, the subjects arrived at the experimental room approximately at noon (**Figure 1B, Nap group**). Then, electrodes were attached for PSG measurement (see ***PSG measurement***) for which it took about an hour. After the electrodes were attached, behavioral task sessions were conducted in the same way as in Experiment 1. After the introductory session in which we explained how to perform the task, the subjects conducted a pre-training test session of the task to measure their initial performance level before training (∼5 min). After the pre-training test session, the subjects underwent intensive training on the task (∼ 45 min). After the training session, the post-training test session (∼5 min) was conducted to measure performance changes by training. There were approximately 2-min short breaks between the pre-training test and training sessions, as well as between the training and post-training test sessions.

Shortly after the completion of the post-training test session, room lights were turned off and the sleep session began at approximately 2 pm, lasting 90 min. This lights-off time being about 2 pm was chosen to take advantage of the effect known as “mid-afternoon dip”, which should facilitate the onset of sleep even in subjects who do not customarily nap (Monk et al., 1996; Horikawa et al., 2013; Tamaki et al., 2016). During the sleep session, PSG was measured (see ***PSG measurement***). Immediately after the sleep session, a questionnaire was administered to collect data about subjects’ introspection regarding the nap (Tamaki et al., 2016). There was a 30 min break after the sleep session to reduce the sleep inertia, which is a prolonged sleepiness upon waking, known to impair performance (Lubin et al., 1976). After the 30-min break, a post-sleep test session (∼ 5 min) was conducted to measure the changes in performance over the sleep session.

Subjective and behavioral sleepiness (SSS (Hoddes et al., 1973), and PVT (Dinges and Powell, 1985), respectively) were measured three times prior to each test session (see ***Sleepiness measurement***).

To collect data about subjects’ sleep habit and handedness, 3 types of questionnaires were administered prior to an adaptation sleep session. They were the Pittsburg Sleep Quality Index questionnaire (PSQI; (Buysse et al., 1989)), the Morningness-Eveningness Questionnaire (MEQ; (Horne and Ostberg, 1976)), and the Edinburgh Handedness Questionnaire (Oldfield, 1971). Using the PSQI, we measured the following parameters: Habitual sleep quality (%), obtained by [(Number of hours slept / Number of hours spent in bed) x 100], the average bedtime, wake-up time, and subjective sleep-onset latency, and the global PSQI score. The global PSQI score indicates the quality of subjects’ habitual sleep (range: 0–21, a global score of >5 suggests poor sleep). The MEQ estimates individual circadian variations. All the subjects in Experiment 2 were right-handed according to the answer in the handedness questionnaire.

### Orientation detection task

A Gabor patch was used for the orientation detection task (**Fig. 1C**). The diameter of the Gabor patch was 10 degrees, presented at the center of the screen. The spatial frequency of the Gabor patch was 1 cycle per degree, and the Gaussian filter sigma was 2.5 degrees. Gabor patches were spatially masked by a noise pattern that was generated from a sinusoidal luminance distribution at a given SNR (Shibata et al., 2017). The average luminance of the stimulus was 130.7± 3.28 cd/m^2^.

Subjects performed the orientation detection task with a two-interval forced choice (2IFC) as in previous studies (Xiao et al., 2008; Shibata et al., 2017). Subjects were presented with two types of displays. One display contained a Gaussian windowed sinusoidal grating (Gabor) patch with a certain signal-to-noise ratio (SNR). The other display had only noise (0% SNR). Each trial started with a 500-ms fixation interval. Two displays were presented sequentially for 92 ms, with a 600-ms inter-stimulus-interval (Xiao et al., 2008; Zhang et al., 2010). After the two displays were presented, subjects were asked to report which display (the first or the second) contained stripes, by pressing the ‘1’ or ‘2’ button on a keyboard.

The threshold SNR was measured for each orientation. The initial SNR was set to 25%. The step size of the staircase was 0.03 log units to adjust the SNR with a 2-down 1-up staircase procedure, which yields about 70% accuracy. The temporal order of the two displays (a Gabor patch + noise or noise alone) was randomly determined in each trial. Subjects were instructed to fixate on a white bull’s eye fixation point (diameter = 1.5 degrees) throughout the display presentations for each trial. No feedback on the accuracy of a response was provided.

The orientation of the Gabor patch was 10°, 70°, or 130°. One orientation was randomly selected as the trained orientation for each subject. Another orientation was randomly selected as an untrained orientation. The remaining one orientation was used for the introductory session (see below). In the training session, a total of 600 trials was performed in 6 blocks (each100 trials) with the trained orientation. The test sessions measured the threshold SNR for the trained and untrained orientations, for each of which one block of staircase was performed. Each block for each orientation ended after 10 reversals, which resulted in about 40 trials per orientation. The geometric mean of the last 6 reversals in each block was obtained as the threshold SNR for the orientation.

The performance improvement (%) by training was calculated as the percent change in the threshold SNR measured at the post-training test session normalized by that at the pre-training test session [performance improvement at post-training: (pre-training – post-training)/pre-training x 100]. The performance improvement (%) by the interval (sleep or wake) was calculated at the post-interval test session relative to the post-training test session [performance improvement at post-interval: (post-training – post-interval)/post-training x 100].

During the introductory session, we explained how to perform the 2IFC Gabor orientation detection task, before the pre-training test in both Experiments 1 and 2. There were approximately 20–30 trials in this session until the subjects reached a certain level of performance. The orientation of the Gabor used for this session was neither trained nor untrained orientation, as mentioned above. In addition, just before the post-interval test session in Experiment 1 and just before the post-sleep session in Experiment 2, the second introductory session was performed with approximately 10 trials to remind the subjects of what the task was.

### PSG measurement

In Experiment 2, the attachment of electrodes for polysomnogram (PSG) measurement, which took approximately 45 min, was conducted prior to the first introductory session. PSG consisted of EEG, electrooculogram (EOG), electromyogram (EMG), and electrocardiogram (ECG). EEG was recorded at 25 scalp sites according to the 10% electrode position (Sharbrough et al., 1991) using active electrodes (actiCap, Brain Products, LLC) with a standard amplifier (BrainAmp Standard, Brain Products, LLC). The online reference was Fz, and it was re-referenced to the average of the left (TP9) and right (TP10) mastoids after the recording. Sampling frequency was 500 Hz. The impedance of active electrodes was kept below 20 kΩ. The active electrodes included a new type of integrated impedance converter, which allowed them to transmit the EEG signal with significantly lower levels of noise than traditional passive electrode systems. The data quality with active electrodes with the impedance below 20 kΩ was as good as 5 kΩ using passive electrodes (Tamaki et al., 2016). The passive electrodes were used for EOG, EMG, and ECG (BrainAmp ExG, Brain Products, LLC). Horizontal EOG was recorded using two electrodes placed at the outer canthi of both eyes. Vertical EOG was measured using 4 electrodes 3 cm above and below both eyes. EMG was recorded from the mentum (chin). The impedance was kept around 10 kΩ for the passive electrodes. Brain Vision Recorder software (Brain Products, LLC) was used for recording. The data was filtered between 0.1 and 100 Hz. PSG was recorded in a soundproof and shielded room.

### Sleep-stage scoring and sleep parameters

Based on the PSG data acquired during Experiment 2, sleep stages were scored for every 30-s epoch, following the standard criteria (Rechtschaffen and Kales, 1968; Iber et al., 2007) into stage wakefulness (stage W), non-rapid eye movement (NREM) stage 1 sleep (stage N1), NREM stage 2 sleep (stage N2), NREM stage 3 sleep (stage N3), and stage REM sleep (REM sleep). Standard sleep parameters were obtained to indicate a general sleep structure for each experiment to confirm that there is no abnormality in subjects’ sleep structures. Sleep parameters included the sleep-onset latency (SOL, the latency to the first appearance of stage N2 from the lights-off), the percentage of each sleep stage, wake time after sleep onset (WASO, the total time of stage W after sleep onset), sleep efficiency (SE, the total percentage spent in sleep), and the time in bed (TIB, the time interval between lights-off and lights-on) (Iber et al., 2007).

### EEG analyses

A fast-Fourier transformation was applied to the EEG data in 5-sec epochs and smoothed with a tapered cosine window (Nobili et al., 1999) to compute brain activities. Six epochs were used to yield the averaged spectral data of 30 s. Spectral powers for sigma activity (13–16 Hz frequency bands) were obtained during NREM sleep stages (from both N2 and N3). Sigma activity of the trained region were obtained by averaging sigma power measured across six occipital electrodes (PO3, PO7, O1, PO4, PO8, O2) that would cover early visual areas, which are assumed to be involved in the Gabor orientation task according to the previous studies (Thut et al., 2006; Viemose et al., 2013). We also obtained sigma activity from the control regions (P7, P8) that are close to the middle temporal gyrus (or MT area) (Seeck et al., 2017). The MT area was chosen as a control, because this region is unlikely to be involved in offline performance gains of a Gabor orientation detection task, given that the MT area is known to be involved in coherent motion perception (Newsome and Pare, 1988; Rees et al., 2000). We computed trained-region specific sigma activity by subtraction of sigma activity of the control region from that of the trained region.

### Statistical analyses

The α level (Type I error rate) of 0.05 was set for all statistical analyses. The Shapiro-Wilk test was conducted for all the data, by which we confirmed that all the data were normally distributed (all *p* > 0.05). The Levene’s test was conducted to test for homogeneity of variance. It was confirmed that homogeneity of variance was not violated for all the data (all *p* > 0.05). The Grubbs’ test was conducted to detect outliers. No outlier data was included in the results.

To analyze performance improvement, ANOVA was first conducted, then *t*-tests were conducted as post-hoc tests. When a correction for multiple comparisons was necessary for multiple t-tests, we controlled the false discovery rate (FDR) (Benjamini and Hochberg, 1995) to be at 0.05. To obtain correlation coefficients, Pearson’s correlation was used.

Statistical tests were conducted by SPSS (ver. 22, IBM Corp.).

## Results

### Experiment 1

We hypothesized that the offline performance gains occur in the Gabor orientation detection task. If this was the case, then the performance improvement over the interval should be larger in the sleep group than the wake group. In addition, since the Gabor orientation detection task has trained-feature specificity (Shibata et al., 2017), we hypothesized that offline performance gains occur with the trained orientation, not with the untrained orientation.

To test whether performance improvement over the interval is larger in the sleep condition, and whether the improvement was specific to the trained orientation, we conducted a 2-way mixed-design ANOVA with Group (sleep vs. wake) and Orientation (trained vs. untrained) factors on the performance improvement at the post-interval test session. The results are shown in **Fig. 2**. The ANOVA indicated a significant 2-way interaction (F(1,6)=6.51, p=0.043), a significant main effect of Group (F(1,6)=12.63, p=0.012) and a significant main effect of Orientation (F(1,6)=16.45, p=0.007). For the trained orientation, the post-hoc tests revealed that there was a significant difference in the performance improvement between the sleep and wake groups but not for the untrained orientation (trained, t(6)=4.11, p=0.006, q<0.05, FDR for 2 comparisons; untrained, t(6)=0.04, p=0.967). Furthermore, one-sample t-tests showed that the performance improvement for the trained orientation at post-interval was significantly different from 0 for the sleep group but not for the wake group (the sleep group, t(3)=7.49, p=0.005, q<0.05, FDR for 4 comparisons including the following 3 one sample t-tests; the wake group, t(3)=0.14, p=0.897). In contrast, the performance improvement for the untrained orientation at post-interval was not significantly different from 0 for both groups (one-sample t-test, sleep group: t(3)=1.74, p=0.181; wake group: t(3)=2.29, p=0.106). Thus, offline performance gains were found for the Gabor orientation detection task, only with the trained orientation. These results were consistent with the hypotheses.

**Fig. 2.**
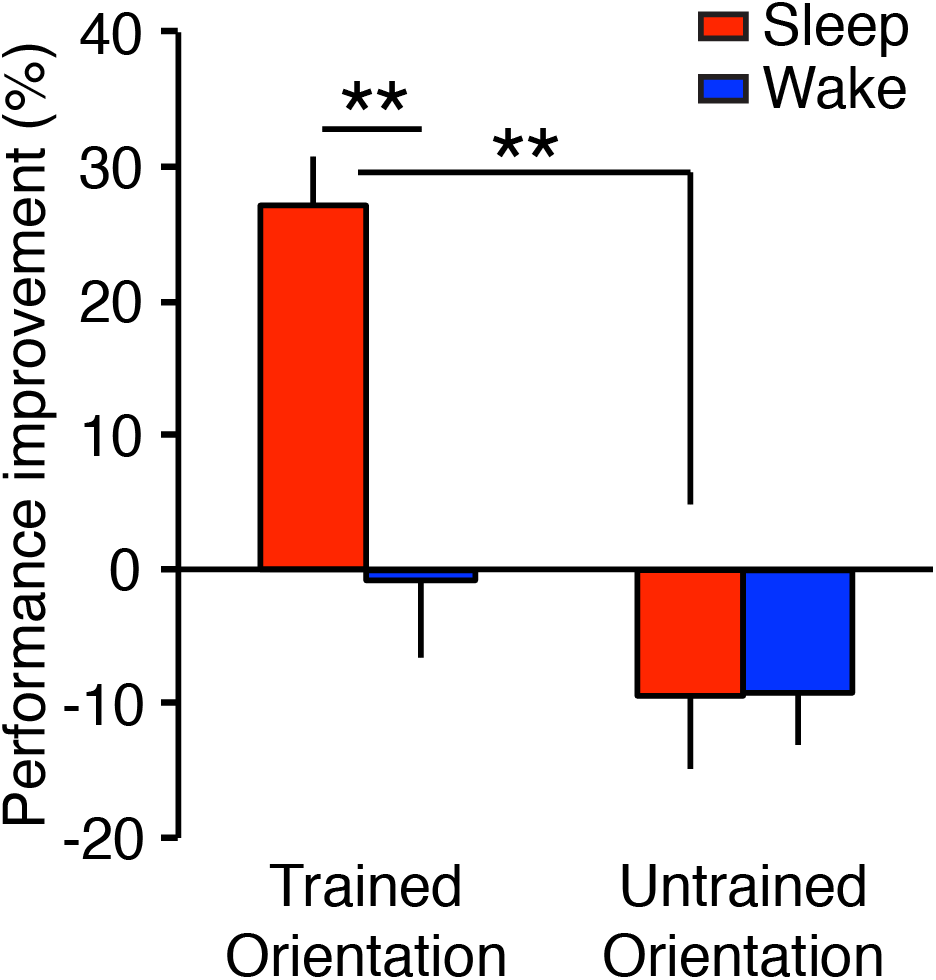
The mean performance improvement (±SEM) at post-interval test session for the sleep and the wake groups in Experiment 1. See the main text for the results of ANOVA. Asterisks (**) indicate that post-hoc t-tests (one sample t-test and a paired t-test) showed significance at p<.01. FDR applied (see the main text for details).

We performed following control analyses to rule out the possibility the difference in the offline performance between the groups was caused by some factors other than the experimental manipulation.

First, we tested whether the initial SNR threshold level was different before training between the groups. However, we did not find a significant difference between the sleep and wake groups in the SNR threshold in the pre-training test session (t(6)=0.07, p=0.945).

Next, we tested whether the training effect was different between the sleep and wake groups. A 2-way mixed model ANOVA with Group (sleep vs wake) and Orientation (trained vs untrained) factors was conducted on the performance improvement at the post-training test session. However, there was no significant main effect of Group (F(1,6)=0.06, p=0.811), no significant main effect of Orientation (F(1,6)=0.02, p=0.899), or no significant interaction between the factors (F(1,6)=0.48, p=0.514) was found. The results indicate that the effect of training was not significantly different between the groups.

Finally, we tested whether the degree of subjective sleepiness was different between the sleep and the wake groups, as the sleepiness may impact on the performance of the detection task. A 2-way mixed design ANOVA with Group (sleep vs wake) and Session (pre-training, post-training, post-interval) was conducted on the SSS ratings. None of the main effect of Group (F(1,6)=0.17, p=0.695), Session (F(2,12)=3.17, p=0.079), or interaction between the factors (F(2,12)=1.50, p=0.262) was significant.

These results indicate that the significant difference between the two groups in the performance improvement over the interval at the post-interval test session cannot be attributed to the initial performance, the effect of training, or subjective sleepiness.

### Experiment 2

In Experiment 2, we tested whether offline performance gains occur with a nap in the Gabor orientation detection task and investigated whether sigma activity during NREM sleep was involved in offline performance gains of the task.

#### VPL performance

We first examined whether the performance on the orientation detection task was improved only in the trained orientation after daytime nap. We conducted one-sample t-tests for the performance changes for each of the trained and untrained orientations at the post-sleep test session. The results indicated that offline performance gains occurred with the nap in the trained orientation (**Fig. 3**). The performance was significantly improved for the trained orientation but not for the untrained orientation (trained, t(8)=3.55, p=0.008, q<0.05, FDR for 2 comparisons; untrained, t(8)=0.25, p=0.811). Furthermore, a paired t-test showed that the performance improvement at post-sleep was significantly different between the trained and untrained orientations (t(8)=2.79, p=0.024). Thus, offline performance gains by nap were specific to the trained orientation.

**Fig. 3.**
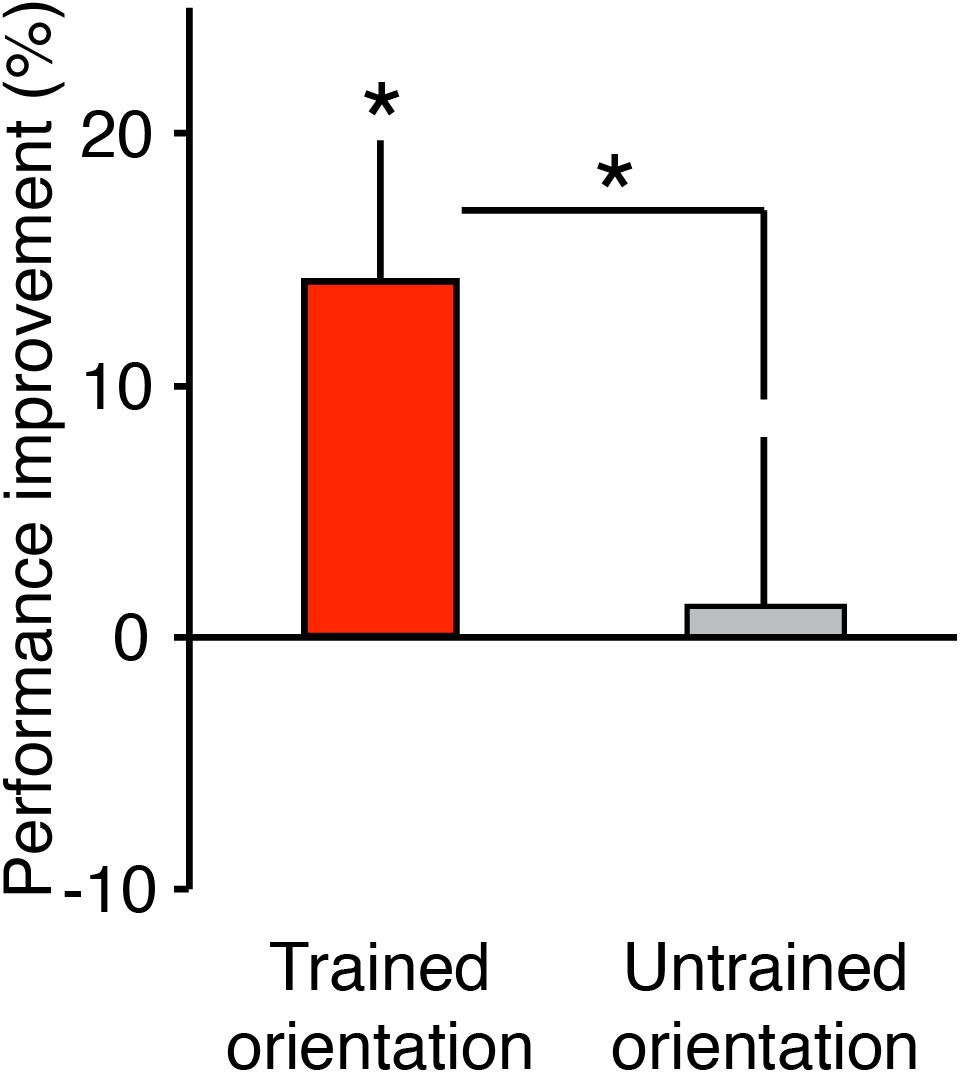
The mean performance improvement (±SEM) at post-interval test session for Experiment 2. N=9. Paired and one-sample t-tests, *p<.05. See main test for more details of statistical results.

Importantly, the difference in the performance improvement between the trained and untrained orientations was not apparent until after sleep. First, there was no significant difference in the initial SNR threshold between the trained vs. untrained orientations at the pre-training test session before training (t(8)=1.29, p=0.234). Second, there was no significant difference in the performance improvement (%) by training at the post-training test session (t(8)=0.65, p=0.531). These indicate that the performance improvement specific to the trained orientation emerged only after sleep.

We compared the amount of offline performance gains obtained in Experiments 1 and 2. There was not a significant difference in the amount of offline performance gains between the experiments (t(15)=0.14, p=0.887), while the amount of the performance gains in Experiment 2 was smaller than that of Experiment 1. Thus, offline performance gains were not significantly different between by night sleep and by a nap.

#### Sigma activity during NREM sleep and the offline performance gain

We tested the hypothesis that sigma activity was involved in the offline performance gain of the detection task in a trained region specific manner. First, we obtained sigma activity in the trained and untrained regions (see ***EEG analyses*** in **Materials and Methods** for more details about regions). We tested whether sigma activity was larger in the trained than untrained regions. We found a significant difference in sigma activity between the regions (paired t-test, t(8)=4.86, p=0.001). Sigma activity was significantly larger in the trained than untrained regions. Next, we tested whether trained-region specific sigma activity was correlated with the performance change over sleep (from the post-training test session to the post-sleep test session, **Fig. 4**, see ***EEG analyses*** in **Materials and Methods**; see **Table 1** for sleep parameters for the sleep session). We found a significant correlation between them (**Fig. 4**, r=0.74, p=0.024, n=9). The Grubbs’ test indicated that there was no outlier in the data. Thus, the result was consistent with the hypothesis that sigma activity was involved in the offline performance gain of the detection task in a trained region specific manner.

**Table 1.** **SOL**, sleep-onset latency. **WASO**, wake time after sleep onset. **SE**, sleep efficiency. **TIB**, the time in bed which indicates the duration of each sleep session (the time interval between lights-off and lights-on)

|  | Mean ± SEM |
| --- | --- |
| Stage W (min) | 21.4 ± 5.61 |
| Stage N1 (min) | 11.2 ± 2.45 |
| Stage N2 (min) | 25.3 ± 4.03 |
| Stage N3 (min) | 18.0 ± 5.30 |
| REM sleep (min) | 5.4 ± 2.61 |
| SOL (min) | 11.0 ± 3.84 |
| WASO (min) | 14.2 ± 4.76 |
| SE (%) | 73.6 ± 7.45 |
| TIB (min) | 85.2 ± 1.96 |

**Fig. 4.**
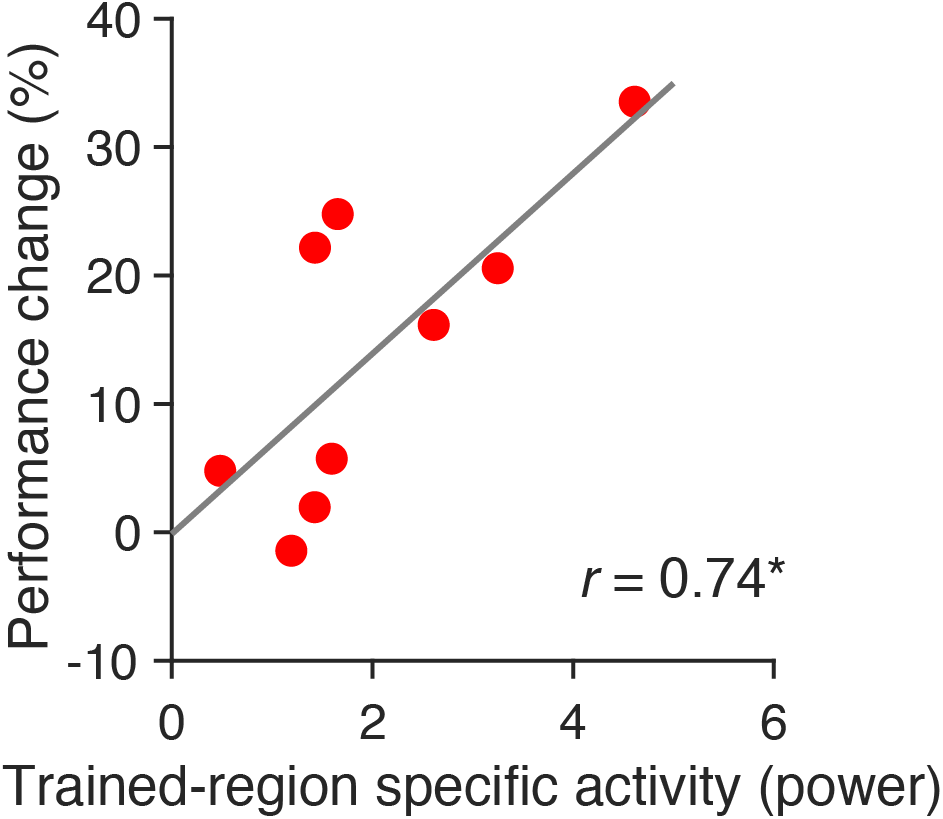
Correlation between the offline performance gains and trained-region specific sigma activity during NREM sleep. N=9. **p<.01. No outliers detected by the Grubb’s test.

We next tested whether any macroscopic sleep variables, such as the duration of each sleep stage was associated with the offline performance gain (see **Table 1** for all the sleep parameters). We measured the Pearson’s correlation coefficient with the performance change over sleep and the durations of each of the sleep stages (W, N1, N2, N3, REM sleep). As a result, none of the durations of sleep stages was significantly correlated with the offline performance gain (stage W, r = –0.18, p=0.642; stage N1, r=0.31, p=0.418, stage N2, r=0.10, p=0.791; stage N3, r = –0.25, p=0.517; REM sleep, r=0.49, p=0.181, without multiple corrections).

We calculated other sleep parameters such as sleep-onset latency (SOL, from the lights-off to sleep onset time (min)), sleep efficiency (SE, the total sleep time divided by the time in bed x 100 (%)), wake time after sleep onset (WASO, min), and total time in bed (TIB (min)) shown in **Table 1**. We tested whether these parameters explain the offline performance gains by calculating correlation coefficients. However, none of them was significantly correlated with the offline performance gains (SOL, r=0.23, p=0.551; SE r=0.18, p=0.645; WASO, r = –0.35, p=0.350; TIB, r=0.17, p=0.669).

#### Quality of habitual sleep and circadian variations

The sleep quality in Experiment 2 was relatively poorer than our recent studies (Bang et al., 2014; Tamaki et al., 2016). The sleep efficiency (%) during the sleep session in the sleep group was 73.6% (**Table 1**). In contrast, in our previous study, the sleep efficiency was about 88.7%-92.8% (Bang et al., 2014; Tamaki et al., 2016) in a sleep session which was done after the adaptation session to eliminate the FNE (Tamaki et al., 2016) (see ***Experimental design for Experiment 2*** in **Materials and Methods** for more information on the FNE). In a similar vein, the WASO in the present study (14.2 min) seems longer compared to the previous studies where it was only 1 – 4.18 min (Bang et al., 2014; Tamaki et al., 2016).

This made us wonder whether the subjects in Experiment 2 were actually poor sleepers, and/or whether they were extreme morning- or evening types. Thus, we analyzed whether the quality of subjects’ habitual sleep was poor using the PSQI questionnaire (Buysse et al., 1989), and whether they were extreme morning or evening type using the MEQ questionnaire (Horne and Ostberg, 1976). The PSQI and MEQ were administered prior to sleep sessions in Experiment 2 (**Table 2**).

**Table 2.** The MEQ score was measured by the Morningness-Eveningness Questionnaire (MEQ). Other parameters were measured using the Pittsburg Sleep Quality Index questionnaire (PSQI).

|  | Mean ± SEM |
| --- | --- |
| Bedtime | 23:16 ± 0:09 |
| Wake-up time | 7:42 ± 0:16 |
| Sleep duration | 7.8 ± 0.20 |
| Sleep onset latency | 11.9 ± 2.57 |
| Habitual sleep efficiency (%) | 92.8 ± 2.25 |
| The global PSQI score | 2.7 ± 0.37 |
| The MEQ score | 55.2 ± 2.63 |

PSQI assesses whether subjects have a sleep problem (Buysse et al., 1989). If the global PSQI score equals or is larger than 5, this suggests that subjects have a sleep problem. However, the average global PSQI score in the subjects in Experiment 2 was 2.7 ± 0.37, ranged 1–4, which indicated that none of subjects was suspected of having sleep problems. In addition, based on the PSQI data, we obtained the subjects’ habitual bedtime, wake-up time, the average sleep-onset latency, the estimated sleep duration, and the habitual sleep efficiency (**Table 2**), These scores in the present subjects in Experiment 2 were considered in normal ranges (Buysse et al., 1989).

MEQ assesses whether the subject is a morning type or evening type (Horne and Ostberg, 1976). Such variations in the circadian timing may affect performance (Kerkhof, 1985). The average MEQ score was 55.2 ± 2.63 with the range between 46–67, which fell into the intermediate type of morningness-eveningness (neither extreme morning nor evening type) (Horne and Ostberg, 1976). These results confirmed that none of the subjects had sleep problems or was of extreme morning or evening type, suggesting that all the subjects had normal sleep-wake habits and were all good sleepers.

We next examined whether any of the measures of sleep habits or habitual sleep quality were related to the performance change. We measured the Pearson’s correlation coefficient between the offline performance gain over sleep and the habitual sleep efficiency, the habitual sleep duration, the global PSQI score, and MEQ score. None of these showed a significant correlation with the offline performance gain (the habitual sleep duration, r=0.54, p=0.134; the habitual sleep efficiency, r=0.52, p=0.150; the global PSQI score, r=0.12, p=0.763; the MEQ score, r = –0.66, p=0.052, without correction for multiple comparisons).

## Discussion

The present results demonstrated that the offline performance gain occurred with a Gabor orientation detection task by nocturnal sleep as well as a daytime nap, in a trained-feature specific manner. Most of the previous studies, which tested offline performance gains in VPL used a discrimination task, in particular, TDT. However, the present study clearly showed that offline performance gains occur with a Gabor orientation detection task. This indicates that offline performance gains are not specific to a discrimination task in VPL.

We found that regional sigma activity in early visual areas was significantly correlated with performance change on the orientation detection task between before and after sleep. It was shown that sigma activity was involved in the offline performance gain of a different task (TDT) over sleep (Bang et al., 2014). Thus, it is possible that offline performance gain is commonly associated with regional sigma activity during sleep in visual areas for VPL.

Sigma activity corresponds to the activities of sleep spindles, which are associated with various types of learning, including learning on declarative and procedural motor tasks (Gais et al., 2002; Schabus et al., 2004; Clemens et al., 2006; Fogel and Smith, 2006; Nishida and Walker, 2007; Tamminen et al., 2010; Tamaki et al., 2013; Laventure et al., 2016; Antony et al., 2018; Boutin et al., 2018). Sleep spindles are specific waves used for sleep stage scoring and appear around the central region during NREM sleep (Rechtschaffen and Kales, 1968; Iber et al., 2007). In the present study, because early visual areas were the targeted regions for offline performance gains of VPL over sleep, we extracted the strength of sigma activity that corresponds to the frequency of sleep spindles from early visual areas. The precise causal relationship between sigma activity and learning are yet to be clarified. However, it has been shown that stimulation whose frequency corresponded to sleep spindles increases long-term potentiation (Rosanova and Ulrich, 2005). Thus, the present results support the idea that regional sigma activity in early visual areas plays a crucial role in offline performance gains of VPL, possibly by increasing regional plasticity in early visual areas during NREM sleep.

The general sleep quality, as indicated in the sleep efficiency and wake time after sleep onset in Experiment 2 seemed to be lower in the present study in comparison to our previous studies (Bang et al., 2014; Tamaki et al., 2016). Although this is beyond the scope of the present study, we speculate that it might be caused by the light exposure from the display of the current stimulus. It has been shown that light exposures before sleep lower the sleep quality and cause more arousals during sleep (Czeisler et al., 1986; Czeisler et al., 1990; Khalsa et al., 2003). As such, it has been shown that playing a computer game with a bright display worsens the sleep quality (Higuchi et al., 2005). In one of our previous studies, which used TDT, the sleep efficiency was about 90% (Bang et al., 2014) while the sleep efficiency was 74.6% in the present study. We measured the luminance for these stimuli. Interestingly, the average luminance for one trial of the Gabor orientation detection task used in the present study was 130.7± 3.28 cd/m^2^, whereas that of TDT was only 0.4 ± 0.07 cd/m^2^. Thus, the current visual stimulus was much brighter than the previous one, as the average luminance of the used visual stimuli was much higher in the present than previous studies (Bang et al., 2014). We speculate that this may be the trigger for the relatively lower quality of sleep in the present study, as other factors, which are known to influence the quality of sleep was controlled and matched between studies including the FNE (Agnew et al., 1966; Carskadon and Dement, 1981; Tamaki et al., 2005; Tamaki et al., 2016), age (Dijk et al., 2000) and habitual sleep problems (Buysse et al., 1989; Breslau et al., 1996; Morin et al., 1999; Morin et al., 2006). In addition, according to the PSQI and MEQ questionnaires, they were of good sleepers and intermediate morningness-eveningness types (Horne and Ostberg, 1976). Thus, only the brightness of the visual stimulus may account for the relatively lower quality of sleep in the present study.

However, it is important to note that the lower quality of sleep was not correlated with performance improvement over sleep in Experiment 2. None of sleep efficiency, wake time after sleep onset, sleep-onset latency and time in bed was correlated with performance improvement. If the wake time during the sleep session was the cause of offline performance gains, there should have been a good correlation for each parameter with offline performance gains. In addition, the results of Experiment 1 clearly showed that the interval that contained sleep led to performance improvements, while the interval that included only wakefulness did not lead to performance improvements. These results altogether demonstrate that offline performance gains of an orientation detection task are a sleep-state dependent process.

In conclusion, the present study demonstrates that offline performance gains were found after training on an orientation detection task. This indicates that offline performance gains are not limited to discrimination tasks including TDT. The present results also suggest that regional sigma activity plays a certain role in facilitation of VPL during sleep.

## Acknowledgements

This work was supported by NIH (R01EY019466, R01EY027841), NSF (BCS 1261765, 1539717), US-Israel Bidirectional foundation (#2016058), and an Institutional Development Award (IDeA) from the National Institute of General Medical Sciences of the National Institutes of Health (P20GM103645). Part of this research was also supported by Center for Vision Research Brown University, and FY17 OVPR Seed Grant, Brown University.

## Author Contributions

M.T. and Y.S. designed the research. M.T. and Z.W. performed the experiments and analyzed the data. M.T., T.W., and Y.S. wrote the manuscript.

## Conflict of Interest

The authors declare no conflict of interest.

